# G protein-Coupled Receptor Distribution Impacts the Effectiveness of Signal Transmission

**DOI:** 10.1101/2020.02.18.953554

**Authors:** Ming-Yi Zhou, Ya-Yu Hu, Huai-Hu Chuang

**Author notes:** To whom correspondence should be addressed: Huai-hu Chuang, Institute of Molecular Biology, Academia Sinica, Taipei, Taiwan 11529. These two authors contribute equally (Ming-Yi Zhou, Ya-Yu Hu).

## Abstract

Numbers of activated receptor dictate efficacy of neurotransmitter stimulation. Many PLC coupled receptors activated by ligands elicit canonical downstream Gq/11 pathway to induce endogenous Ca^2+^ gated chloride channels. The coupling from receptors to effectors was analyzed in *Xenopus* oocytes expressing genetically modified angiotensin receptor type 1 receptor (AT1R). The latency between ATII binding and Ca^2+^-induced Cl^−^ current surge was inversely correlated. AT1R activation triggered a chain of chemical reactions, of which the products were playing messengers for subsequent events. Messenger accumulation must rate-limit the agonism. For accurate quantification the speed of ATII triggered the *i* Cl^−^. The T-form AT1R-IRK1 fusion exhibits faster induction compared to the M-form. The latency of the recorded none vanished *i* Cl^−^, marking the lowest genuine calcium activation, took place at earlier time point by the timer time. The evoked *i* Cl^−^ however reached similar maximal amplitudes. This kinetic effect raises the possibility to use temporal coding to complement amplitude coding (analogous to FM versus AM radio transmission) for receptor-agonist pairs.

## Introduction

There are several families of transmembrane proteins encoded by almost a thousand genes that are critical for normal physiology, with G protein-coupled receptors (GPCRs) being the largest and most ubiquitous [1]. The highly specific ligand–GPCR interaction prompts an efficient cellular response, which is vital for the health of cells and organisms. GPCRs share seven conserved membrane-spanning alpha-helical transmembrane domains that are activated by ligand binding in the extracellular space [2, 3]. Through conformational changes, ligand-bound GPCRs activate heterotrimeric G-proteins, which execute the downstream signaling pathways through recruitment and activation of cellular enzymes [4].

Up to 40% of drugs target GPCRs [5]. Despite considerable progress in using GPCRs as drug targets, our understanding of how these drugs act on GPCRs remains incomplete. GPCRs can be activated by different ligands to transduce similar signaling pathways. Previous studies employing various techniques including fluorescence microscopy, crystallography or theoretical modeling have revealed aspects of GPCR signaling transduction at the molecular level [6-11]. However, since the receptor-transducer complex is highly dynamic and can adopt many conformational transitions, it is virtually impossible to use crystallographic techniques alone to capture all conformational states of either bound or unbound GPCRs. Moreover, the resolution of conventional methods is insufficiently sensitive to quantify small changes in receptor number. Time-resolved measurements are essential to characterize the order of events leading to formation of GPCR-transducer complexes, especially when paired with complementary functional studies.

Angiotensin II type 1 receptor (AT1R) belongs to a subtype of GPCRs [12, 13] and plays an critical role in modulating hypertension [14]. AT1R has been identified in a wide variety of tissues including the kidney, liver, adrenal gland, cardiovascular system, and brain [15-17]. Upon binding to angiotensin II (ATII), AT1R is stabilized in its active conformation and stimulates heterotrimeric G proteins. G proteins dissociate into alpha (Gq/11 family) and beta/gamma subunits. AT1R interacts primarily with Gq/11 proteins in many ATII target cells [18, 19]. Gq/11 subunits act as signal transducers to activate phospholipase C (PLC), leading to hydrolysis of phosphatidylinositol 4,5-bisphosphate (PIP_2_) and formation of diacylglycerol (DAG) and inositol trisphosphate (IP_3_). DAG and IP3 stimulate the endoplasmic reticulum (ER) to release intracellular Ca^2+^ [20-25].

In the brain, many hormones and neurotransmitters activate cellular signal transduction pathways via G-protein coupled receptors [26, 27]. Gα proteins are speedily activated upon the formation of ternary complexes from direct interaction of receptors with their cognate ligands. Among a collection of PLC-coupled GPCR pathways, despite all signaling through the canonical Gαq and downstream effectors thereof, each receptor still elicits events of a distinct spatiotemporal profile. To elucidate the basis on which such versatility arises, we performed single cell analysis to monitor GPCR activation in real time.

Receptor-effector coupling is an under-studied step in signal transmission. The analytical process is inherently complex given that relevant factors determining the outputs in either upstream or downstream end are usually multifactorial so that the outputs have a wide dynamic range [28-32]. Such data scattering obscures the underlying mapping from inputs to outputs. To address the basis of coupling efficiency, the derived quantity from both ends of the equation must be accurately measured; analysis of coupling also demands a range of inputs evenly distributed in the spatial as well as time domain in the broadest technique-possible range, especially for deriving the threshold of signal induction at which the receptor number would expectedly be so low as to hamper its precise measurement. Therefore, we use sensitive and precise electrophysiological methods to uncover the usually accompanied low-level expression.

*Xenopu*s oocytes are a widely used model for studying heterologous expressed receptors, transport proteins, and ion channels. To answer the coupling mechanism by which GPCRs transduce downstream signaling, the high level of endogenous Ca^2+^ activated Cl channels (CaCCs) could be measured as a reporter. CaCCs are activated by elevated intracellular Ca^2+^ concentrations regardless the Ca^2+^ source. When the Ca^2+^ concentration reaches a sufficiently high threshold, the chloride channel opens to allow significant Cl^−^ efflux. Thus, GPCRs induce signaling events to convey a rapid sub-plasma membranous Ca^2+^ concentration rise, which could be detected precisely by the conducted Cl^−^ current through CaCC.

We considered that the numbers of AT1R receptor dedicate the rate of the ATII-induced chloride currents responses; more AT1R receptor numbers would have a faster response rate. In this study, we selected inwardly rectifying K^+^ channel (IRK1), an ion channel insensitive to most known cytosolic messengers and cellular modulators [33-36], to construct a receptor-channel fusion. We found that this method can accurately calculate AT1R receptor numbers by measuring potassium flux through recombinant receptors expressed in *Xenopus laevis* oocytes. With agonists bound to the AT1R receptors, the resulting activated G proteins induced downstream pathways by elevating intracellular Ca^2+^ concentrations to induce Cl^−^ flux, allowing us to investigate the relationship between receptor numbers and the rate of chloride current production.

## Results

### Expression of M-form and T-form of AT1R-IRK1 fusions in *Xenopus* oocytes

We developed a technique to indirectly infer the number of G-protein receptors expressed by detecting channel currents, and to quantify the efficiency of the receptors. By generating fusions of receptors and channels together, the ion channel activity could serve as the readout for receptor activation. We have designed the pGEMHE vector with elements that enable efficient protein expression in the *Xenopus* oocytes (Figure 1A). A fused sequence with the AT1R gene linked to an inwardly rectifying potassium channel K_ir_2.1 (IRK1) gene, which was engineered into the pGEMHE to generate AT1R-IRK1 fusion proteins (Figure 1B). We constructed two types of fusion proteins, M-form and T-form of AT1R-IRK1. The M-form fusion protein includes an AT1R fused with an IRK1 subunit, whereas the T-form fusion protein has an AT1R linked to a tetrameric tandem string of IRK1 subunits. Since four IRK1 subunits form a functional tetrameric K channel [41], the M-form of AT1R-IRK1 represents a functional IRK1 channel linked four AT1R receptors and the T-form of AT1R-IRK1 represents a functional IRK1 channel linked an AT1R receptor.

**Figure 1.**
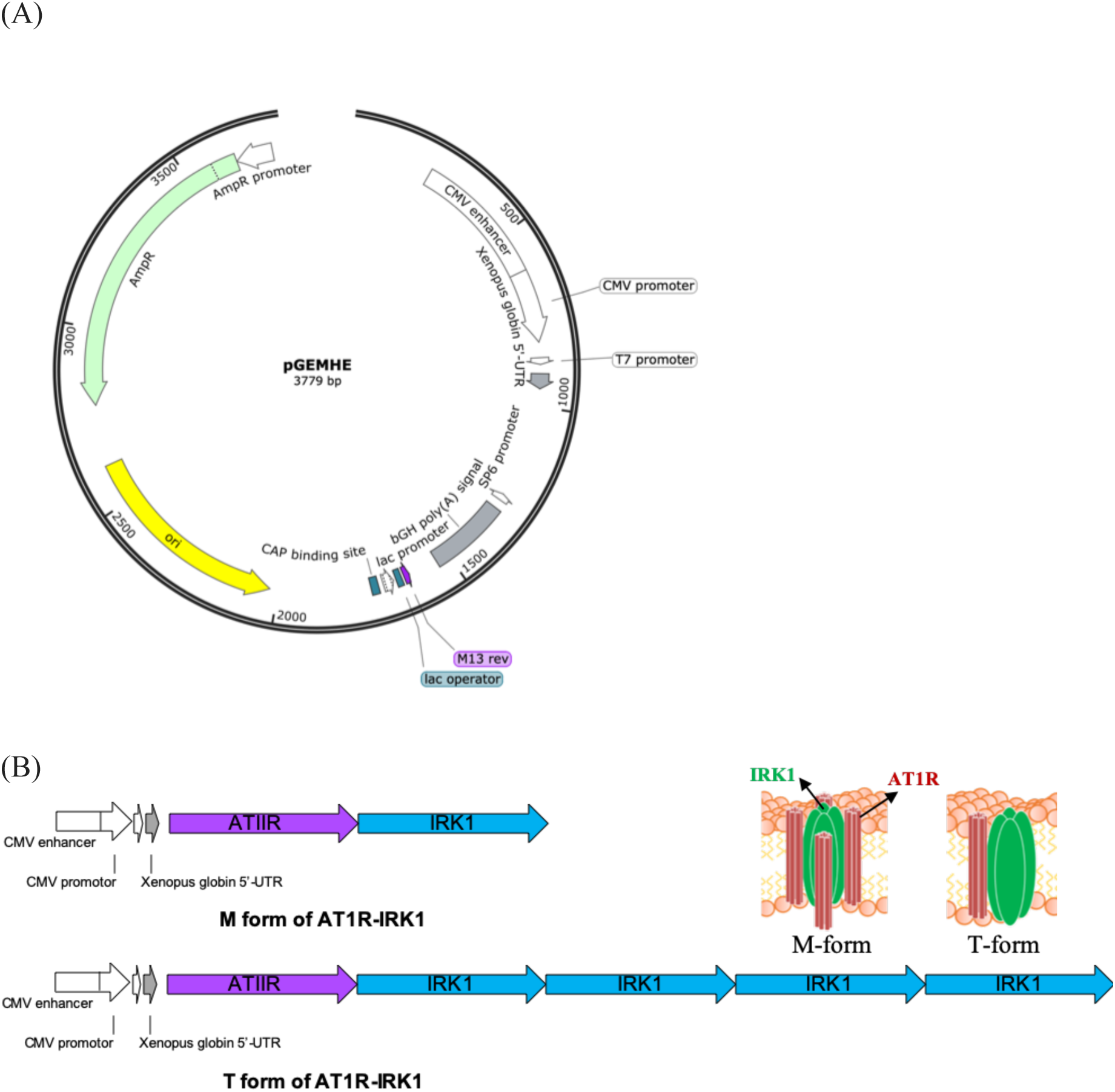
AT1R-IRK1 construction. (A) The pXpIV vector contains an origin of replication (*ori*) (*yellow segment*) and an ampicillin resistance gene (*Amp*^*r*^) (*Green segment*) for maintenance in ampicillin-sensitive bacterial strains. Expression in mammalian cells is driven by the CMV immediate early enhancer/promoter, and cRNA transcription uses T7 RNA polymerase. The *Xenopus* globin 5’-UTR contains the bovine growth hormone polyadenylation signal (bGH poly(A) signal). The *Xenopus* globin 5’-UTR and the polyA tail including the bovine growth hormone polyadenylation signal were included to improve the stability of RNA in oocytes and to boost the protein expression level [39][40]. The chimeric intron is ATII linked with IRK1, which be inserted between the CMV promoter and the bGH poly(A) signal. (B) Construction of AT1R-IRK1, M-form and T-form. The M-form of AT1R-IRK1 includes one ATIIR gene followed with an IRK1 gene. The T-form of AT1R-IRK1 is one ATIIR gene followed with four IRK1 gene. Schematic diagram of proposed M-form and T-form of AT1R linked with and IRK1 structure.

### Functional IRK1 in M-form and T-form of AT1R-IRK1

We utilized the two-electrode voltage-clamp technique to record potassium currents by Ramp program (Figure 2A). To confirm that the function of IRK1 channels is not be altered, we examined the IRK1 currents in oocytes expressing M-form or T-form of AT1R-IRK1 with the voltages from −120 to +80 mV in 600 ms. We showed that the AT1R-IRK1 fusion proteins expressed heterologously in *Xenopus* oocytes with biophysical hallmarks of IRK1, including high K^+^ selectivity and strong rectification. The current-voltage (IV) curves of the M-form and the T-form AT1R-IRK1 showed no difference to IRK1 alone (Figure 2B). These results indicate that IRK1 is functional when fused to the AT1R, and the recorded K^+^ currents might be suitable for measuring the numbers of AT1R.

**Figure 2.**
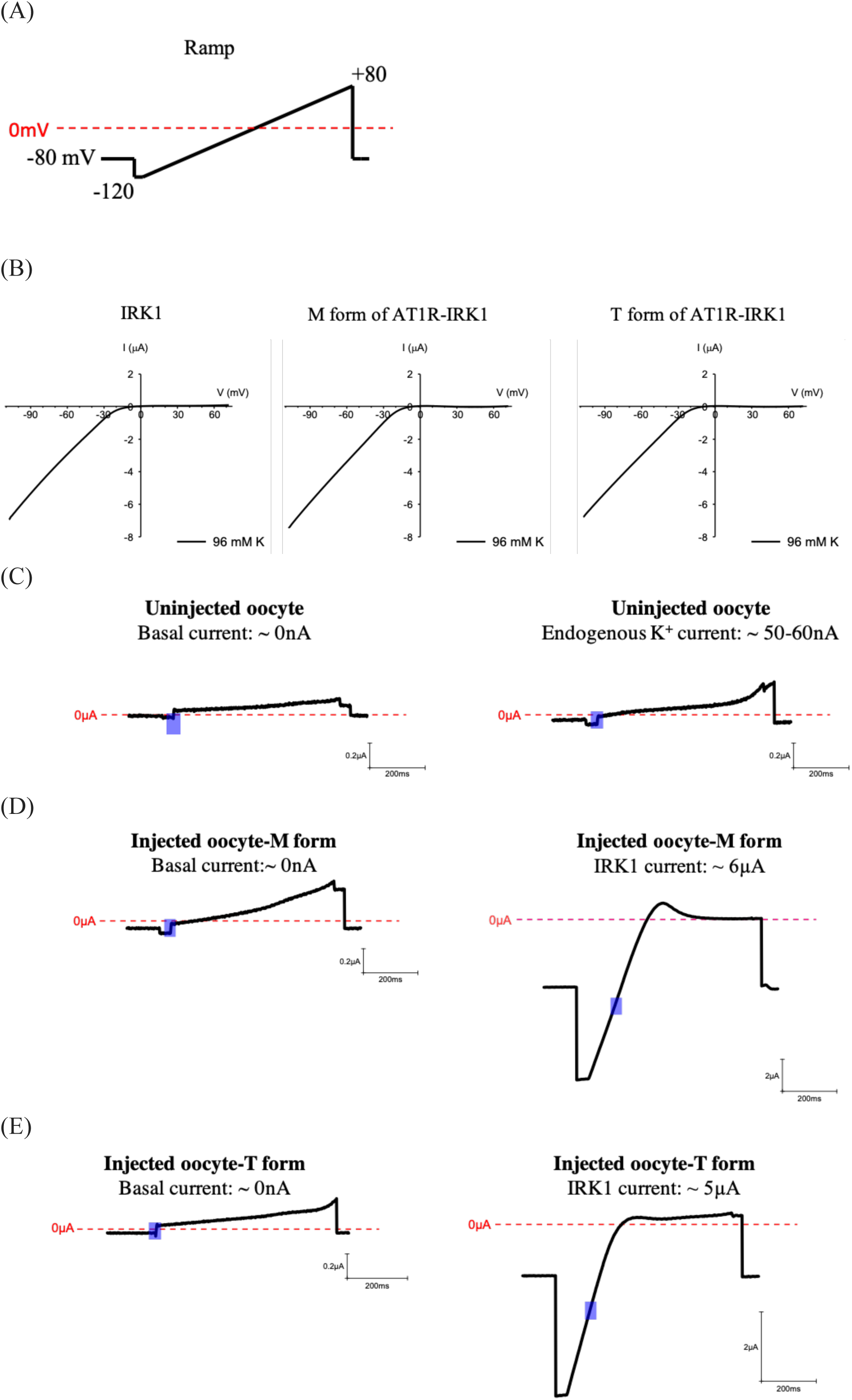
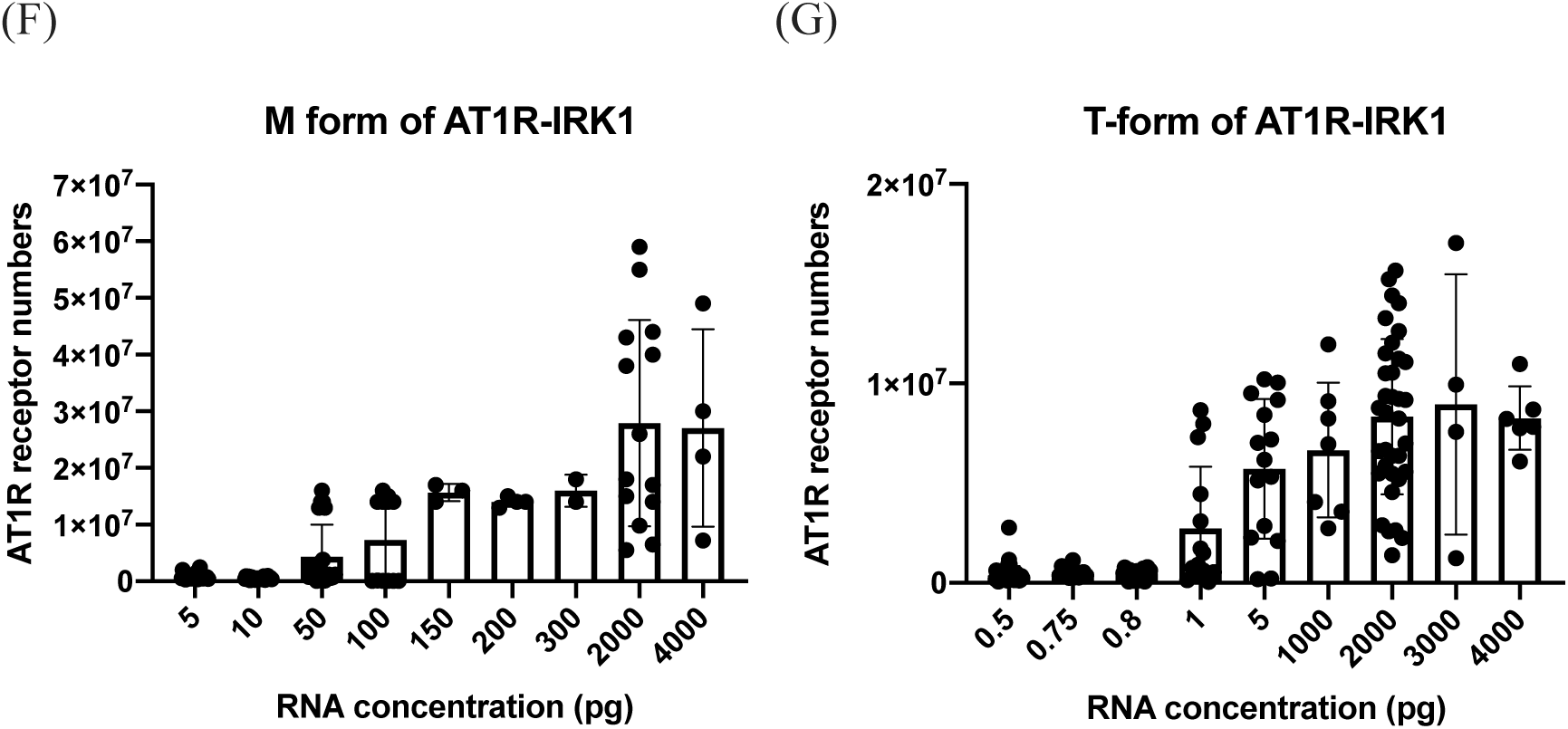
Quantitative analysis of AT1R numbers by recording IRK1 current utilizing two-electrode voltage clamp. (A) We designed a Ramp stimulus program with the voltage change from holding potential −80mV switched to −120mV at 160ms, increasing from −120mV to +80mV in 600ms, and from +80mV switched to the holding potential at −80mV at 860ms. We measured the IRK1 current with the average voltage change from −114mV to −94mV, and we recorded the chloride current with the voltage holding at +80mV. (B) The I-V curves showed that IRK1 current alteration with voltage change. We compared the IRK1 current in three group, IRK1 alone, M-form of AT1R-IRK1 and T-form of AT1R-IRK1. We injected 500 picogram RNA of M-form and T-form in *Xenopus* oocytes. All of them are perfused with 96mM potassium solution. (C) Uninjected RNA oocytes were used to be a comparing group as injected oocytes. We firstly measured the basal current, nearly close to 0nA at normal Na^+^ (96 mM) extracellular solution without pottasium ion in uninjected oocytes, then we switched to normal K ^+^ (96 mM) extracellular solution to record their endogenous potassium current, around 50 to 60nA with the average voltage change from −114mV to −94mV (shown in purple box). (D) The oocytes expressed the M-form of AT1R-IRK1were measured the basal current 0nA at normal Na^+^ (96 mM) extracellular solution and IRK1 current 6μA in K ^+^ (96 mM) extracellular solution with the average voltage change from −114mV to −94mV (shown in purple box). (E) The oocytes expressed the T-form of AT1R-IRK1were measured the basal current, nearly close to 0nA in Na^+^ (96 mM) extracellular solution and IRK1 current 5μA in K ^+^ (96 mM) extracellular solution with the average voltage change from −114mV to −94mV (shown in purple box). (F, G) We injected different concentration of M-form and T-form of AT1R-IRK1 RNA into oocytes. The graph shows the relationship between RNA concentration and AT1R receptor numbers.

While *Xenopus* oocytes are an outstanding heterologous expression system for investigating ion channel activity, the oocytes express an amazing variety of endogenous potassium ion channels that can interfere with electrophysiological measurements. Therefore, we recorded the endogenous potassium current in uninjected oocytes. In the K^+^-free, 96 mM NaCl perfusion solution, the basal current was nearly 0 nA (Figure 2C). In the 96 mM K^+^ solution, the oocyte endogenous potassium currents were around 50 to 60 nA in the voltage range of −94 mV to −114 mV. These recorded potassium currents in uninjected oocytes were used as basal control. Then we examined the IRK1 currents in oocytes injected with RNA of M-form or T-form AT1R-IRK1. Both basal currents recorded from M-form and T-form in the 96 mM NaCl solution showed no difference to the basal current recorded from the uninjected oocytes (Figures 2D and 2E, left panels). After perfusing with 96 mM K^+^ solution, the injected M-form RNA or T-form RNA showed a significant IRK1 current, respectively (Figures 2D and 2E, right panels). These results indicate that the IRK1 channels in M-form and T-form AT1R-IRK1s are functional.

### Quantifying AT1R numbers from recorded potassium currents

Next, we tried to use the IRK1 currents to measure numbers of AT1R. Previous study showed that IRK1 single channel conductance is around 21pS (Aleksandrov A., et al. 1996) at 150 mM K^+^ solution, which was employed as IRK1 single conductance in the calculation. We measured the conductance stable change (slop) region of recorded current trace (at −114 ∼ −94 mV, purple boxes in Figures 2C-E). The average conductance change from ten repeated recordings was further divided by the value of IRK1 single-channel conductance. The values of calculation showed the numbers of IRK1 expressed at the oocyte surface, which means numbers of AT1R as well. Both M-form and T-form of AT1Rs were obtained in increasing amounts of AT1R-IRK1 fusion RNAs injected into oocytes, thereby allowing us to establish how differential expression of fusion proteins drove the observed phenotypes.

Next, we examined the relationship between the concentration of injected RNA and numbers of AT1R-IRK1 fusions. With increasing RNA concentrations, both receptor numbers of M-form and T-form AT1R-IRK1 increased accordingly. In M-form, the amounts of RNA ranging from 5 to 4,000 pg were injected. We found that in some cases the maximum receptor number could reach as many as 60 millions. The average maximum receptor number is at about 30 millions with injection of 2,000 pg of RNA, which could not be further boosted by increasing the amount of injected RNA to 4,000 pg. Also, the minimal RNA required to elicit significant response is 50 pg (Figure 3A). In T-form, the RNA amounts ranging from 0.5 pg to 4,000 pg were injected. Interestingly, 1 pg of RNA was sufficient to elicit K^+^ current for measurement, and the average maximum receptor number is at 10 millions with injection of 2,000 pg RNA. Independent maximum receptor numbers could reach as high as 20 millions (Figure 3B). These data showed that the minimal and maximal receptor numbers expressed in oocytes between M-form and T-form are different. The difference in the M-form and T-form in converting injected RNA into the oocytes to the measured active receptor numbers on the surface could be several folds (see Discussion). We believe that the measured K^+^ current better reflects the number of active receptors on the surface than the amount of injected RNA, which are used for further study.

**Figure 3.**
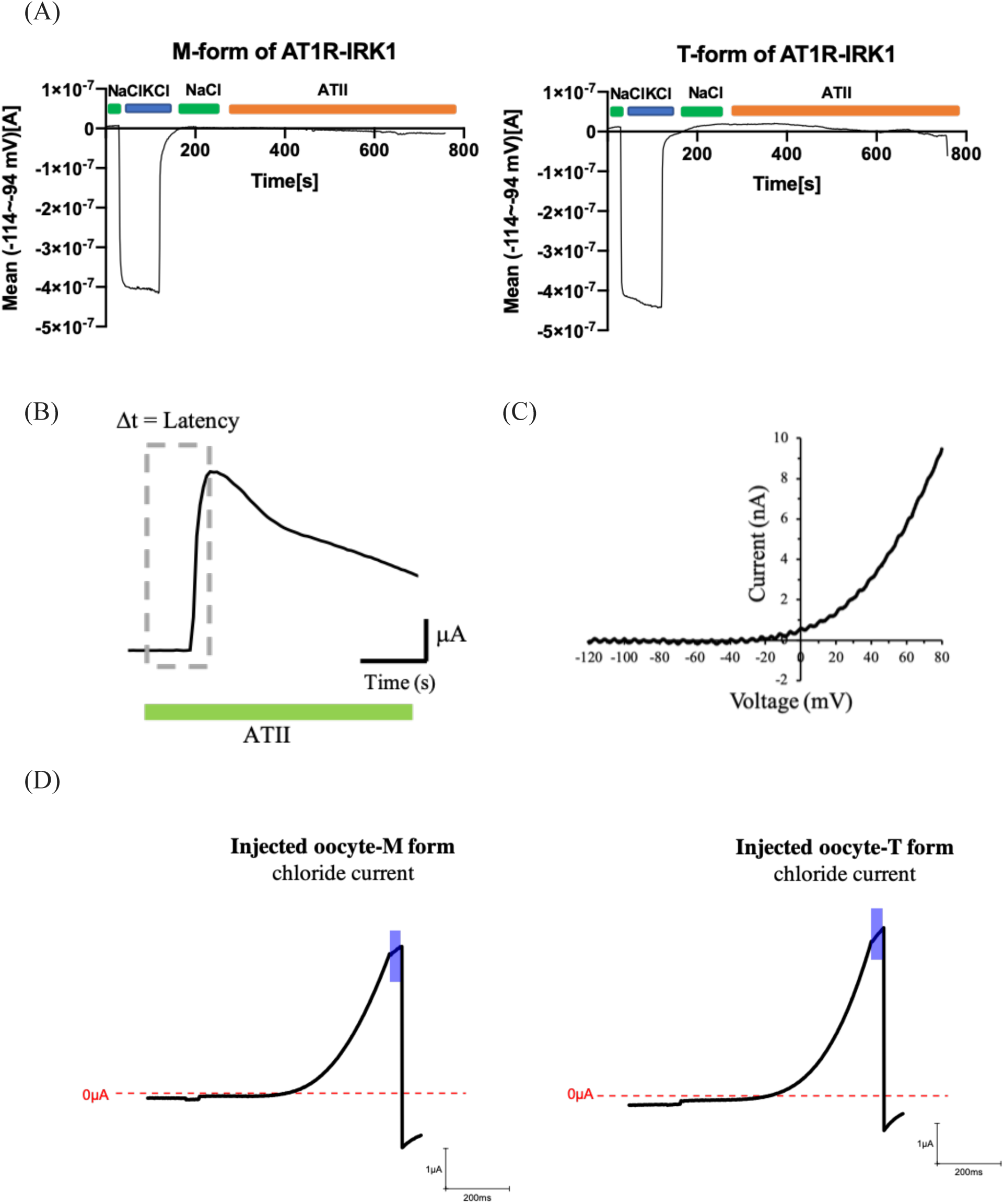
ATII-induced calcium-activated chloride current in M-form and T-form of AT1R-IRK1 receptor. (A) The Ramp Program was set with 610 ramp within around 14 minutes. *Xenopus* oocytes injected with RNA of M-form or T-form initially perfused with 96mM NaCl solution without potassium ion from Ramp 1 to 28, swithced 96mM KCl soultion from Ramp 29 to 106, switched to 96mM NaCl solution from Ramp 107 to 235, and finally perfused with 1μM AT1R recptor agonist 1 μM angiotensin II (ATII) from Ramp 237 to the end. (B) The latency was difined as depicted, the time from administering 1μM ATII to induce the first chloride current response. (C) The IV curve indicated the chloride current in the Ramp program at +80mV. (D) The oocytes expressing 500pg RNA of M-form and T-form of AT1R-IRK1were measured the chloride current administered 1 μM ATII in 96 mM NaCl solution. The curves are ATII-induced chloride current of M-form and T-form. The purple area indicates the average voltage (+59mV to+79mV) of measuring chloride current.

### ATII induced calcium-activated chloride currents in AT1R-IRK1 receptors

To understand the efficiency of response rate, we measured the activity of AT1R receptor in the presence of the agonist angiotensin II (ATII). When ATII binds to AT1R receptor, active G-protein generates a cascade of downstream signaling transduction, inducing (CaCC) open and Cl^−^ efflux. To calculate the correlation between the number of AT1R and the maximum Cl^−^ current of CaCC, we established a continue Ramp with totally 610 repeats. Both M-form and T-form were tested (Figure 3A). Because the rate of signal transduction downstream in the whole cell is constant, the time from the activation of AT1R to the peak chloride current (latency, Figure 3B) should be proportional to the amount of total activated AT1R on the cell surface. With the addition of agonist ATII, the first peak of the chloride current was recorded within 640 ms (ramps) from +59 mV to +79 mV of Ramp (Figure 3C). The basal current obtained from ten consecutive points prior to the application of ATII was subtracted from the measured chloride current to establish the activated chloride current.

Next, we examined whether the function of AT1R functions would be altered. While the K^+^ current was recorded, the Cl^−^ current was also measured in both M-form and T-form upon the ATII application. The detection of Cl^−^ currents in both AT1R-IRK1 fusions suggests that function of AT1R normally in both M-form and T-form fusions (Figure 3D).

### T-form of AT1R-IRK1 exhibits higher chloride current induction efficiency than M-form of AT1R-IRK1

The receptors of the cells appear randomly and uniformly, and the individual responses to stimuli should be consistent. We investigate whether receptor distribution affects the efficiency of whole cells response. We plotted the response rate of ATII-induced Cl^−^ current against the number of AT1R-IRK1 fusions. With low number of M-form receptors (∼ a few millions), the response rate was very low. With increasing receptors, the response rate increased linearly and reached plateau when the receptor number reaching over 40 millions (Figure 4A). This result indicated that the response rate of M-form fusion has the maximum close to the 0.2 s^−1^. Interestingly, the T-form fusion response rate also increased quickly and semi-linearly even in the range of a few millions receptor, reaching 0.2 s^−1^ with fewer than 20 millions (Figure 4B). Therefore, the response rate on the number of receptors for binding to the agonist ATII indicates that T-form AT1R-IRK1 has a higher signal transmission rate than that of M-form AT1R-IRK1.

**Figure 4.**
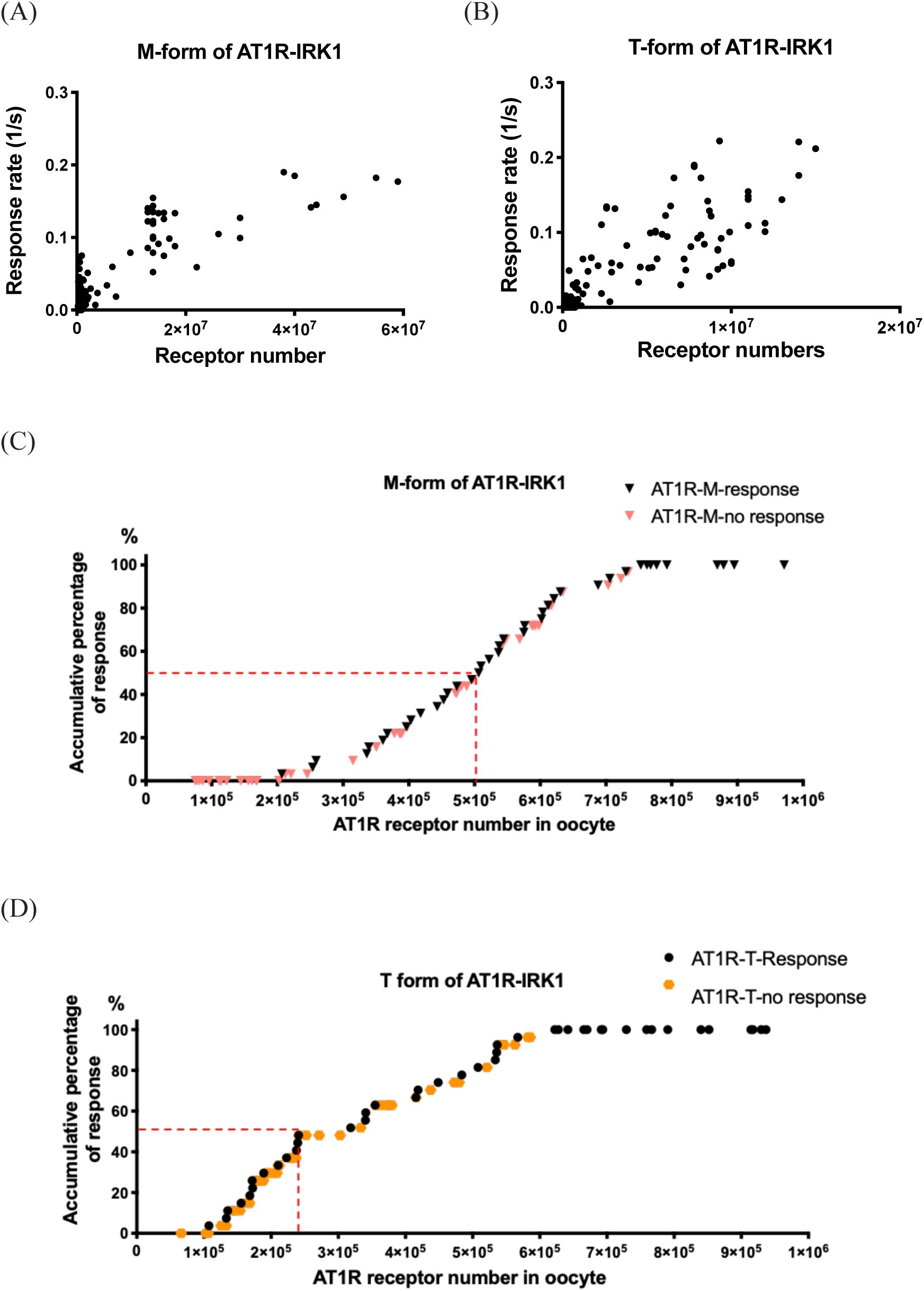
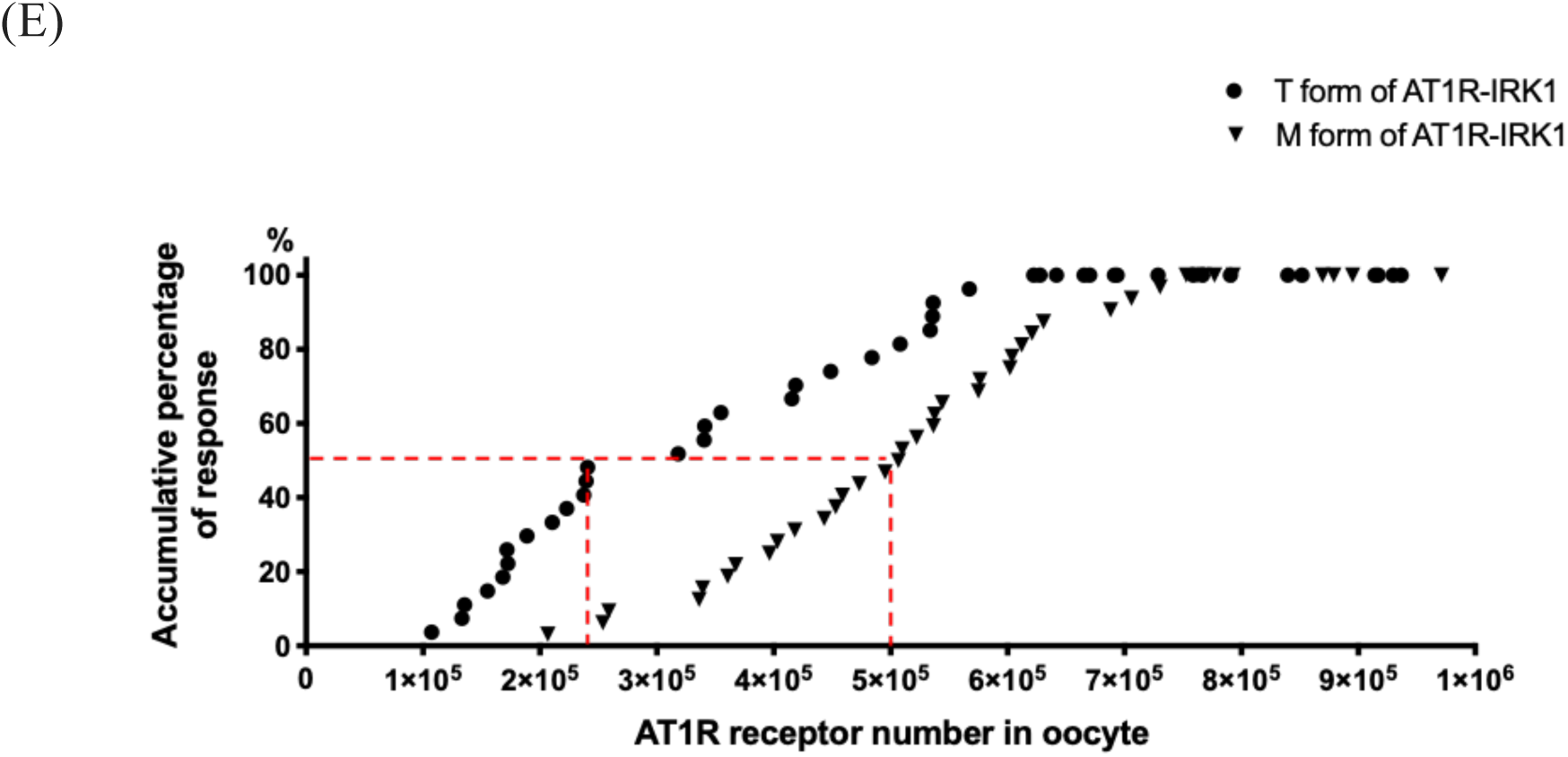
Fewer activated T-form AT1R-IRK1 receptors than M-form are required to induce the first calcium-activated chloride channel (CaCC) response. (A) The relationships between the oocyte response rate upon administration of 1 μM angiotensin II and AT1R receptor numbers in our two recombinant AT1R-IRK1 fusion proteins, M-form and T-form. (T-form of AT1R-IRK1, n = 141; M-form of AT1R-IRK1, n = 103) (B)(C) The thresholds of upstream triggers depend on the intrinsic threshold for Ca^2+^ activation of a CaCC response. The probability of production chloride current in M-form and T-form of AT1R-IRK1 Each point represents an oocyte. Black group represents that ATII binds AT1R receptors and produces the chloride current, while orange group represents that ATII binds AT1R receptors but without producing the chloride current. (D) The relationship between the number of AT1R-IRK1 fusion proteins and the cumulative percentage of induced CaCC current recorded for intervals of one hundred thousand receptors. More than seven hundred thousand AT1R fusion proteins are required to induce a maximal cellular response. The threshold number of M-form AT1R-IRK1 is about five hundred thousand, while the threshold number of T-form AT1R-IRK1 is about two hundred fifty thousand.

We assessed the minimum number of receptors required to transduce extracellular signals into oocyte responses. Our use of Ca^2+^-activated Cl^−^ current as a powerful reporter considerably enhanced the sensitivity of our assay. We plotted a cumulative probability distribution curve against the number of receptors within 1 million (Figure 4C, 4D). The M-form fusion showed that there was no detectable Cl^−^ current production below 2 × 10^5^ receptors. Even administrating the higher concentration of ATII, undetectable Cl^−^ current response was produced (data not show). Between 2-7 × 10^5^ receptors, stochastic responses were still detected. With more than 7 × 10^5^ receptors, consistent responses were ensured in each RNA injection. The T-form fusion showed that detectable Cl^−^ current was produced even at 1 × 10^5^ receptors, and consistent responses were detected with more than 6 × 10^5^ receptors. For T-form fusion proteins, 50% successful ATII-induced stimulation occurs when fewer than 3 × 10^5^ receptors were present or, to be more accurately, agonist-bound receptors are present on an oocyte, whereas for M-form receptors, that number is 5 × 10^5^ receptors (Figure 4D). The results show that M-form and T-form fusion receptors show different response probability and cell sensitivity.

Thus, physiologically, ATII must act on a sufficient number of receptors on a cell membrane to trigger downstream cell responses. Thus, when an oocyte has up to ∼ 7 × 10^5^ agonist-bound receptors on an oocyte, the T-form has a higher probability of inducing Cl^−^ current responses than the M-form, a surprisingly novel result.

## Discussion

In this study, the AT1R was made into a channel fusion to facilitate synchronous receptor counting and concurrent monitoring of ligand-induced response. We could quantitatively the response for AT1R according to the receptor numbers of each. Thanks to the selectivity of potassium channels, we could separate the K-currents from a mixture of currents by simple ion replacement. We deduced the receptor number by measuring the whole cell current in K^+^ containing solutions, subtracting the background non-K current in the K^+^ free condition. The net K currents were converted into the number of receptors by dividing to the single channel conductance. This measurement is simple, direct, and stoichiometrically proportional. We set up a wide dynamic range of receptor expression, granting the subsequent quantitative analysis for coupling. The question we asked first was how many receptors would a cell need in order to initiate a whole-cell agonist response, i.e., the neurotransmitter induced calcium-activated chloride current?

Multiple studies have demonstrated that many GPCRs exist as dimers or high oligomers [42, 43]. Some of these studies have also shown that GPCR dimerization or oligomerization is important for receptor function, including in relation to agonist affinity, potency, and efficacy, as well as G protein specificity [44-48]. Moreover, upon ligands binding to receptors, activated G proteins induce conformational changes of the receptors that may influence the function of other proximal receptors. Thus, the distance between receptors may alter their functions [49-51].

We created the one-to-one receptor-channel fusion. The receptor portion of the fusion was directly ligated to a tandem tetrameric IRK1, T-form of AT1R-IRK1. By this design, no local high receptor density would be possible. If expressed at low enough level, one would not anticipate the receptors formed on the cell surface capable of assembling into local complexes. With the limitation of receptor numbers in oocytes membrane, the fusion of an IRK1 subunit by design must use their hydrophobic surface to assemble into tetrameric channels (a 4-receptor-to-1 receptor-channel design), M-form of AT1R-IRK1, which should bring the ligand binding sites physically closer than the 1-to-1 isomers in limitation of spatial distribution in oocytes membrane. However, someone might suspect that the creation of linked tandem receptor had created the higher-order receptor tetramers. With four AT1Rs fused to an IRK1 channel possibly caused unintentional and unnatural alteration of the receptor activity. Surprisingly, these two different designs did not display any measurable functional differences in receptor-effector coupling.

However, cell membrane package of M-form and T-form receptors could be very different. Our examination of chloride current response latencies reveals that, to exert the same oocyte activation rate, fewer T-form agonist-bound receptors are required than M-form receptors. Reduced the latency does not necessarily mean a stronger chloride current response. There are some exceptions that short latency was accompanied with smaller Cl^−^ current response. But by and large, short latency was associated with a larger Cl^−^ current which fired more readily. It is possible that quarternary structures of the M-form are more constrained than for the T-form. The physical distances between agonist (ATII) binding sites in oocytes harboring M-form receptors may be closer than for those hosting the T-form because formation of a channel pore in the former necessitates proximity of four IRK1 subunits whereas, in the latter, each agonist binding site is independent.

The spatial distribution of AT1R receptor raises the possibility of using temporal coding to complement amplitude coding for receptor-agonist pairs. This situation is likely to FM versus AM radio transmission. Adequate receptor numbers located at the appropriate sites can produce responses, while insufficient or superabundant receptor numbers, or their locations too crowded or too loose may not able to induce the responses. In our study indicated that at the same receptor numbers, T-form of AT1R-IRK1 have rapid Cl^−^ current response rate than M-form of AT1R-IRK1. This phenomenon suggested that AT1R receptor sites of M-form are so crowded that each receptor coupled G protein influences each other to their downstream signaling pathway.

The receptor-channel fusion had been a convenient and effective intermediate tool for counting the number of the receptors. The inherent risk of building a complex not already existing in nature is failure to form the intended structure because of folding problems ending in low yield, insolubility or accelerated degradation. We bypassed this stage and got the designated proteins made in the Xenopus oocytes. The engineered protein expressed sufficiently and folded properly and targeted to the expected destination. In the functional analysis, they behaved similar to their native counterparts.

Receptor-effector coupling depends upon every element involved in between [52]. To obtain a response, receptor number must reach a threshold to set off the cascade activation of CaCC. The timing of the CaCC chloride current surge upon receptor activation, i.e. latency of the response, is a measurable indicator with which we can analyze the coupling efficiency between the upstream events and the effectors responses, ligand exposure on the surface and the calcium store release and any other responses following respectively. Typically, a period of membrane quiescence named latency was noted between exposure to soluble ligands from external leaflet and a rise of Cl current amplitude peaking shortly. Albeit the current amplitudes widely variable, latencies remain simple and robust for easy recording and subsequent detailed analysis.

When ATII binds to AT1R, not only may it activate Gq protein to induce the IP_3_ downstream pathway but it can activate another G protein, Gi, to induce the cAMP-dependent pathway [53-55]. If Gq protein levels are too low to induce sufficient calcium release from the ER, CaCCs may not be activated, thereby preventing the generation of chloride current. Differential ligand binding affinities may also explain the variation in chloride current elicited by T-form and M-form receptors. Due to conformational changes in receptors upon ligand binding, G protein coupled efficiency may be potentially different for T-form or M-form receptors.

The acceleration of CaCC activation presents itself as a reduction of latency. Spatial expansion of the signal-transducer-acting domain should endow an increase of downstream response amplitudes or reduction of failure rates along coupling. We systematically varied the receptor numbers or altered the arrangement of relative spatial positions for agonist binding. This analysis is more sensitive and of higher precision than the direct measurement of delay, promising the potential of receptor activity for more incisive modeling. It opens the gate to revisit GPCR action wanting more general and useful parametric values in quantitative analyses, for instance, partial agonism or spared receptors.

In our study, we measured the time required between binding of angiotensin and the onset of chloride currents, latency. It is a reproducible receptor-ligand pair specific parameter. The application of ATII to a cell expressing ATIIR-fusion activated Gaq, hence the downstream chloride currents, which always display a waiting period with electrical quiescence. It reflected the delay to the time-dependently accumulate second messengers Ca^2+^. The latency was rate-limited by the production of GTP bound Gaq before the system reached the saturation expression of receptor. Expectedly, the latency was inversely correlated to the receptor expression level. It therefore indicated how effective the receptor activation was translated into an ionic response.

In sum, the AT1R-IRK1 fusion expressed as a full length polypeptide folded properly for both moieties; it gave rise to a strongly inwardly rectifying potassium channel with an unchanged single channel conductance; the receptor portion behaved like the parental GPCR with the identical ligand recognition; above all, activation by cognate agonists were of the same specificity and affinity. Application of neurotransmitters set off the coupling to the chloride channel CaCC. The more the K currents were the speedier the onset of CaCC, shown from the shortened latency. Reciprocals of latencies indicated how rapidly the effector response starts. Provided that receptor numbers were sufficiently low, the latency would be long enough to make confident measurements revealing the efficiency of receptor-effector coupling. We chose latency measure instead of other parameters particularly the size of CaCC to score post-receptor events to avoid over-amplification of downstream response thereby weakened quantitative causal relationship. The shortened latencies indicated an increase of speed of cytosolic calcium rise, a consequence of the IP_3_ surge. A larger post-receptor response arises from multiple steps of amplification. Although all three receptors we chose for latency analyses coupled to the same pathway, their coupling efficiencies differed. Importantly, the thresholds of upstream triggers depended on the intrinsic threshold for Ca activation of CaCC. By contrast, the relative efficacy of the GPCR receptor was determined as the time passed before achieving a certain percentage of maximal Gaq responses.

If the initial step dictates most of the final outcomes, one might wonder why so many intermediate steps exist between the trigger and the effector. One potential advantage of setting up many intermediate events is that they could serve as buffering steps, either by sequestering the mediating elements or by augmenting replenishment of those components lost to attrition, ensuring the success of downstream transduction. In all, we illustrates that the robustness of the signaling system relies critically on the driving power of the trigger, which is a gate to control whether a response can be elicited. However, the overall resolution resides in the late steps on the effector end. Our analysis in this case is generally applicable, given that principles of GPCR transduction mechanisms are highly conserved across different subclasses of G-proteins. Such a conceptual framework will underlie the quantitative basis for the robustness of GPCR signaling.

## Materials and Methods

### AT1R-IRK1 fusion protein construction and heterologous expression of fusion protein

The constructs used in this study included angiotensin II type I receptor (AT1R) (human) and Kir2.1 (GIRK, rat). Complementary DNAs were subcloned into plasmid vector pGEMHE designed for expression in *Xenopus* oocytes. Plasmids were then linearized with *Smil* and capped RNA was transcribed with T7 transcription kit (mMessage mMachine; Ambion, Austin, TX). The yield and quality of transcripts was assessed by agarose gel electrophoresis. Stages VI oocytes from *Xenopus tropicalis* anesthetized by 0.4% ethyl 3-aminobenzoate methanesulfonate (MS-222)(Sigma-Aldrich; St. Louis, Missouri, USA) were prepared by treatment with 0.1 mg/ml collagenase (Worthington, type CLS3) for 30 minutes at room temperature with agitation in 82.5 mM NaCl, 2 mM KCl, 1 mM MgCl_2_, 5 mM Hepes (pH 7.4). Separated and defolliculated oocytes were then rinsed and stored in 82.5 mM NaCl, 2 mM KCl, 1 mM MgCl_2_, and 5 mM Hepes (pH 7.4) at 18°C with 100 μg/ml streptomycin and 100U/ml penicillin. Up to 24 hr after collagenase treatment oocytes were injected (Nanoject; Drummond, Broomall, PA) with cRNA in 18.4 nl volumes of different concentrations. For some experiments, it was necessary to inject more RNA to produce functional effects; we have specified the quantities of RNA used in these instances.

### Electrophysiology

All experiments were conducts at room temperature (22°C). For two-electrode voltage clamp experiments, the potassium-free extracellular solution contained 96 mM NaCl, 1 mM MgCl_2,_ 1 mM CaCl_2,_ 5 mM HEPES (PH7.4), from which 96 mM Na^+^ was replaced with K^+^ to make the standard extracellular K^+^(96K) solution. Recording typically started in K^+^-free solution and was switched to 96K solution to obtain basal potassium currents. Angiotensin II (ATII) was applied in the 96 mM Na^+^ solution. The oocyte membrane potential was held at −80 mV and given a voltage ramp from −120 mV to +80 mV of 600-ms duration applied at 1Hz. The IRK1 exhibits a significant potassium current flow when the membrane potential is clamped at −120 mV. Utilizing recording K^+^ current subtracted basal current and then divided by a functional IRK1 conductance obtained the numbers of ITK1 channel. Then we used the IRK1 channels numbers to calculate AT1R receptor numbers.

When ATII were administered to fusion channel-protein, activated G proteins trigger a series of downstream pathways that cause fluctuations in cytosolic calcium concentration were ultimately inducing CaCC current responses. We used the maximum concentration 1μM angiotensin II as an agonist and measured the average chloride peak current (from +59 to +79 mV) of the recorded trace in the RAMP electrical stimulation mode. We can monitor the experimental time variation of this average current. Owing to a series of downstream signal pathways are considered to be the same single response, the time between the administration of the agonist and the induction of the first Cl^−^ current is referred to as the latency. The Cl^−^ current response induction rate is the reciprocal of the latency. Data are acquired and analyzed with the PatchMaster software (HEKA). Data are analyzed and presented by origin (Northampton, Massachusetts, USA).

## Author contributions

HHC and MYC designed the experiments, MYC and YYH performed the molecular and electrophysiological experiments. YYH integrated the experimental results. YYH and HHC wrote the manuscript. HHC supervised their progression.

## Acknowledgements

We thank Dr. Cheng-Ting Chien for helping this manuscript completed. We also thank Dr. Hsueh-Chi Yen, Dr. Yi-Fang Tsay and Dr. Jun-Yi Leu for stimulating discussion and insightful comments on the manuscript.

